# Pairwise and higher-order epistatic effects among somatic cancer mutations across oncogenesis

**DOI:** 10.1101/2022.01.20.477132

**Authors:** Jorge A. Alfaro-Murillo, Jeffrey P. Townsend

## Abstract

Cancer occurs as a consequence of multiple somatic mutations that lead to uncontrolled cell growth. Mutual exclusivity and co-occurrence of mutations imply—but do not prove—that they can exert synergistic or antagonistic epistatic effects on oncogenesis. Knowledge of these interactions, and the consequent trajectories of mutation and selection that lead to cancer has been a longstanding goal within the cancer research community. Recent research has revealed mutation rates and scaled selection coefficients for specific recurrent variants across many cancer types. However, estimation of pairwise and higher-order effects—essential to estimation of the trajectory of likely cancer genotoypes—has been a challenge. Therefore, we have developed a continuous-time Markov chain model that enables the estimation of mutation origination and fixation (flux), dependent on somatic cancer genotype. Coupling the continuous-time Markov chain model with a deconvolution approach provides estimates of underlying mutation rates and selection across the trajectory of oncogenesis. We demonstrate computation of fluxes and selection coefficients in a somatic evolutionary model for the four most frequently variant driver genes (*TP53, LRP1B, KRAS* and *STK11*) from 565 cases of lung adenocarcinoma. Our analysis reveals multiple antagonistic epistatic effects that reduce the possible routes of oncogenesis, and inform cancer research regarding viable trajectories of somatic evolution whose progression could be forestalled by precision medicine. Synergistic epistatic effects are also identified, most notably in the somatic genotype *TP53*+*LRP1B* for mutations in the *KRAS* gene, and in somatic genotypes containing *KRAS* or *TP53* mutations for mutations in the *STK11* gene. Large positive fluxes of *KRAS* variants were driven by large selection coefficients, whereas the flux toward *LRP1B* mutations was substantially aided by a large mutation rate for this gene. The approach enables inference of the most likely routes of site-specific variant evolution and estimation of the strength of selection operating on each step along the route, a key component of what we need to know to develop and implement personalized cancer therapies.

## 1 Introduction

Abnormal cell proliferation and survival can be driven by gene mutations in somatic cells, and can result in cancer. The somatic mutations that lead to cancer can become frequent within tumors either because they are frequently mutated or because they are strongly selected to increase proliferation and survival. However, the selection operating on somatic mutations is complicated by epistatic effects, wherein the effects of one mutation affect the selection operating on other genes [1].

Mathematical models in oncology can help to understand the mechanisms of oncogenesis, cancer progression, and optimal treatment [2]. Computational models have revealed extensive pairwise epistasis among germline variants and among gene knockdowns [3, 4], and approaches have been applied to identify sets of variants that are sufficient to cause cancer [5]. Among somatic mutations, patterns of mutual exclusivity have been interpreted as a consequence of antagonistic epistasis, and patterns of co-mutation have been interpreted as a consequence of synergistic epistasis. Indeed, pairwise epistatic interactions can vary with genomic background [6], and substantial higher-order epistasis is common in evolutionary genetics [7]. Therefore, to reveal the evolutionary genetic trajectories of cancer that could be intercepted by precision therapeutic approaches to delay or deny cancer morbidity and mortality due to metastasis, quantification of higher order epistatic effects are needed [8]. In cancer, higher-order epistatic models proposed so far do not acknowledge the extensive gene- and site-specific variation in underlying mutation rates [9], nor condition on them or their correlations [10]; nor do they reveal the order of mutations revealed by the epistatic interactions. For example, a mutation in a gene *A* could make it more likely that a mutation in a gene *B* is acquired, whereas when the mutation in the gene *B* occurs first, the mutation in gene *A* could be selected against [11]. These differences in selection can arise from stage-specific physiological divergence or from additional unrecognized epistatic interactions. The estimation of the order of epistatic effects in cancer is particularly challenging because most large tumor sequence datasets provide only one time point of tumor evolution at the time of tumor biopsy or excision [12]. Most evaluate correlations between the frequencies of mutations [3], yet correlations can arise either because of selective epistasis or because of commonalities of mutation process between selected sites. There are several orders of magnitude of difference in the rates at which somatic mutations occur in different genes and sites within the genome. Therefore, it is vital to distinguish the mutation rate from how much one mutation would confer a selective advantage in the tumor cell population [9]. The scaled selection coefficient or cancer effect size has previously been averaged over epistatic effects [9]. Evaluating epistatic effects of selected mutations in cancer evolution is fundamental to illuminating the evolutionary genetic trajectory of tumor evolution, and for the advancement of accurate and personalized predictions of therapeutic responses [13–15].

Herein, we develop a mathematical model enabling the inference of the potential evolutionary trajectories of tumors. Our approach extends from pairwise epistasis to combinations of three or more cancer drivers. We apply our models to lung cancer, the most frequent cancer in the world [16], and the leading cause of cancer death in the United States [17]. Lung adenocarcinoma represents about 40% of all lung cancers, and has the worst prognosis among all types of lung cancer [18]. It typically has a high tumor mutational burden [19], and thus provides an excellent example in which to test theory on epistatic effects in cancer oncogenesis.

## 2 Theory

### 2.1 Mutational flux for one mutation with no epistasis

For a single mutation, *λ* can be taken as the exponentially distributed flux from a “normal” tissue state (without a mutation) at time 0 to a tissue state with a mutation fixed throughout a neoplasm, then

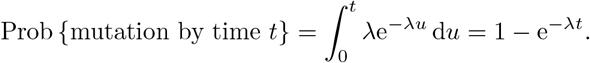

This flux can be decomposed into the neutral mutation rate (the rate at which the genetic state is changed in single cells and would get fixed without any selective pressure) times the scaled selection coefficient (the consequent change in survival and proliferation due to selection) [9].

### 2.2 Mutational fluxes for two mutations with epistasis

For two mutations labeled *A* and *B*, the fluxes from a normal state to a state with mutation *A* and to a state with mutation *B* can be denoted *λ*_*A*_ and *λ*_*B*_. Assuming a regime of strong selection and weak mutation (SSWM) [20] with no clonal interference on the action of selection, and thereby retaining exponential distributions for the time that it takes for each mutation to appear and be selected to high frequency, the minimum of those distributions is also exponential, with the exponential parameter equal to the sum of the parameters of each distribution. Starting in the normal state at time 0, the probability density function for the time until the first mutation fixes is

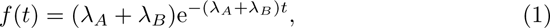

for *t* ≥ 0 and 0 otherwise. Therefore, the probability of maintaining the normal tissue state through time *t* is

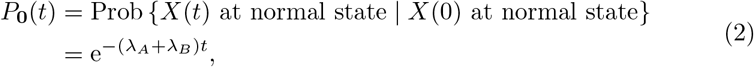

for *t* ≥ 0, where *X*(*t*) represents the state at time *t*. Additionally, we know that under SSWM, the probability that a specific mutation spreads to fixation before another mutation is equal to its rate relative to the sum of event rates, thus

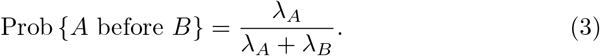

Without epistasis, the fixation probabilities of mutation *B* are independent of whether mutation *A* has previously risen to fixation in the tumor. Therefore under SSWM the probabilities of fixing either mutation or both can be computed by multiplying the respective probabilities for each event. However, widespread mutual exclusivity among driver mutations indicates that epistatic interactions between them may be commonplace [16, 21, 22]. To model epistatic interactions between two driver mutations, two additional parameters are required: the flux to mutation *A* while at a state with mutation *B*, denoted *λ*_*B*→*AB*_, and the flux to *B* while at *A*, denoted *λ*_*A*→*AB*_. Thus, for a tissue fixed for mutation *A* at time *u* and *t* ≥ *u*,

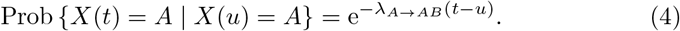

The probability of a tissue fixed with only mutation *A* at time *t* can be computed by multiplying Equations (1), (3), and (4), and integrating:

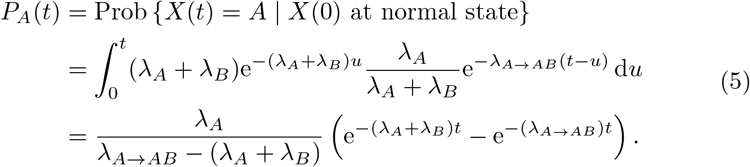

Equations (1), (3), and (4) compose three conditions that together yield the desired probability that a tissue fixed only mutation *A* at time *t* (Equation(5)). Equation (1) conditions that a mutation fixed at a time *u*, Equation (3) conditions for the case that the mutation was *A* instead of *B*, and Equation (4) conditions for the case that no other mutations were fixed from the time *u* that the mutation was fixed to the time *t*.

By symmetry,

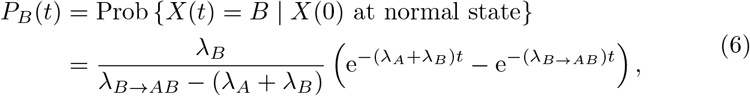

and a formula for the probability that the neoplasm will be in a state fixed for both mutations follows from Equations (2), (5), and (6):

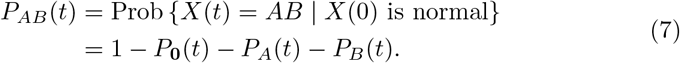

For two mutations, Equations (2), (5), (6), and (7) provide probabilities for all possible genotypic states of the evolving neoplasm.

If three or more mutations are considered, an equation for the probability of having two mutations fixed cannot be obtained by subtracting the other probabilities as in Equation (7). Alternatively, *P*_*AB*_(*t*) can be directly computed with the formula

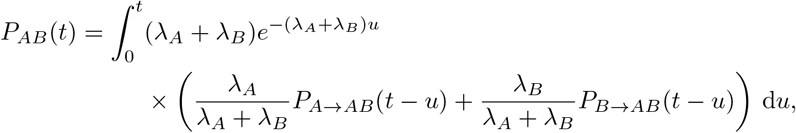

where

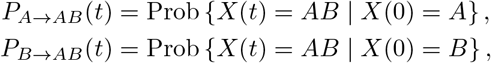

are probabilities that can be obtained by a similar argument as the one used to obtain Equation (5). However, as we only require *P*_**0**_(*t*), *P*_*A*_(*t*), *P*_*B*_(*t*) and *P*_*AB*_(*t*) for the likelihood (Section 3.2), it would be better to obtain a formula where *P*_*AB*_(*t*) is in terms of *P*_*A*_(*t*) and *P*_*B*_(*t*). We will obtain such an equation for the general case with *M* mutations.

### 2.3 Mutational fluxes for *M* mutations with epistasis

To solve the general case of *M* possible somatic mutations, we can model the somatic genetic state of a neoplasm with respect to time *t* as a continuous-time Markov chain *X*(*t*), *t* ≥ 0. We define the set of all possible states of the system as the *M* -ary Cartesian product S = {0, 1}^*M*^. Any state in the system is represented by a vector in S:

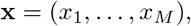

where *x*_*i*_ is 1 if the state has fixed the *i*-th mutation and 0 otherwise. Modeling two mutations *A* and *B, M* = 2; the normal state would be represented by (0, 0), the state with only mutation *A* fixed by (1, 0), the state with only *B* fixed by (0, 1), and the state with both *A* and *B* fixed (*AB*) with (1, 1).

Under an SSWM regime, mutations occur and spread one at a time. Consequently, the flux from **x** to **y** is 0 unless **y** has exactly one more mutation than **x**, or **y** is **x**. Thus, the infinitesimal parameters *λ*_**x**→**y**_ that determine the flux from state **x** to **y** are such that *λ*_**x**→**y**_ = 0 unless there is an *i* ∈ {1, …, *M*} with *x*_*i*_ = 0, *y*_*i*_ = 1 and *x*_*j*_ = *y*_*j*_ for all *j* ≠ *i*, or **y** = **x**. The **y** = **x** case is relevant because—as is customary for continuous-time Markov chains—we define

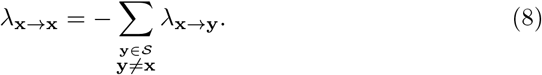

If **x** ≠ (1, …, 1), Equation (8) can be simplified to

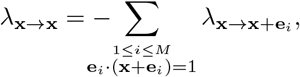

where **e**_*i*_ represents the *i*-th vector of the standard basis of ℝ^*M*^, that is, the vector with zeros everywhere except for a 1 in its *i*-th entry. The state with all mutations fixed **x** = (1, …, 1) is an absorbing state, therefore the infinitesimal departure rate from the state is zero, i.e. *λ*_(1,…,1)→(1,…,1)_ = 0.

Modeling *M* = 2 mutations, all infinitesimal parameters can be represented by the continuous Markov matrix:

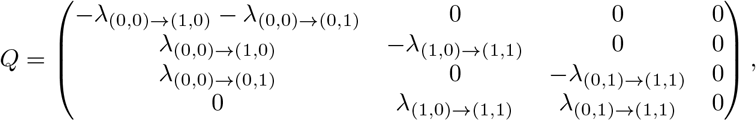

with the ordered rows and columns representing the states (0, 0), (1, 0), (0, 1) and (1, 1). Equivalently, this matrix can be written with the notation of Section 2.2 as:

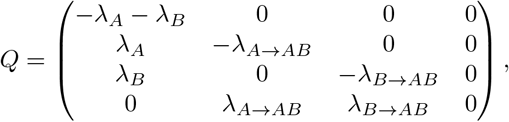

with ordered rows and columns representing tissue in a normal state, a state with only the first mutation *A*, a state with only the second mutation *B*, and a state with both mutations *AB*.

Because *X*(*t*) is a continuous-time Markov chain, the probabilities that a neoplasm starts in a state **y** at time *u* and is in a state **x** at time *t* + *u* are independent of *u*, so we can denote them as:

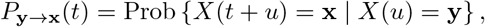

for any *t, u* ≥ 0. Applying Kolmogorov’s backward equation [23] to the matrix *P* (*t*) with entries 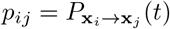, for any ordering 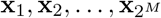 of all possible states, we obtain the differential equation

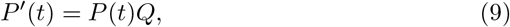

where *Q* is the continuous Markov matrix for the same ordering of all possible states. Thus we could find the solution for *P* (*t*) by computing the matrix exponential for *Q* and applying the Fundamental Theorem for Linear Systems [24,Section 1.4], then:

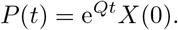

If all eigenvalues of the matrix *Q* are real and distinct then the exponential matrix can be found by finding its diagonalization [24, Section 1.2]. Performing this operation and solving for the probabilities of being in the mutation states associated with *M* = 2 (0, *A, B*, and *AB*) yields Equations (2), (5), (6), and (7). However, a direct solution for the exponential matrix is exhaustive to compute for large matrices [24, Section 1.8]. The size of the matrix *Q* grows exponentially with *M*. Even for the case of *M* = 3 an 8 × 8 matrix is required, and each entry of *e*^*Q*^ is prohibitively complicated. To develop an alternate approach toward quantification of the probabilities of state for *M* mutations, a main property of our estimation problem may be capitalized upon: it can be assumed that all individuals start at the normal state, that is, without relevant somatic mutations. Therefore, only one column in the matrix *P* is of interest, the one that includes the entries *P*_**0**→**x**_. To simplify notation, we will write

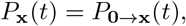

for any *t* ≥ 0. The relevant column in Equation (9) reduces to

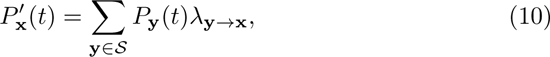

where the specific ordering of state transitions is no longer important as it was in Equation (9). By the properties of the continuous-time Markov chain, we know the initial condition for this differential equation:

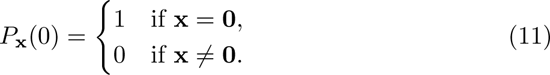

For the case of **x** = **0** = (0, …, 0), that is, the normal tissue state, Equation (10) becomes

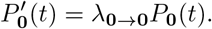

Solving this differential equation with the initial condition specified by Equation (11), we have

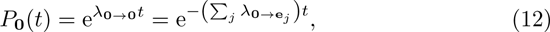

where the sum goes for all *j* = 1, …, *M*.

For the case that **x** ≠ **0**, using Equation (10) and the definition of the infinitesimal parameters,

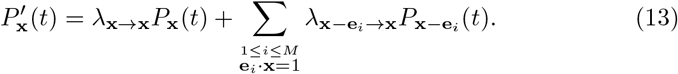

Solving Equation (13) with the integral factor 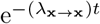 and the initial condition in Equation (11) provides a recursive formula

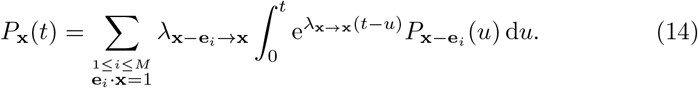

This formula enables computation of *P*_**x**_(*t*) for all states **x** by starting with the normal state using Equation (12), then proceeding to compute all states with one mutation (where Equation (14) depends on *P*_**0**_(*t*)), then all states with two mutations (where Equation (14) depends on *P*_**x**_(*t*) for states **x** with only one mutation), and so on.

A straightforward validation case of the Equation (14) comes from its application when **x** already features exactly one mutation. Applied to that case,

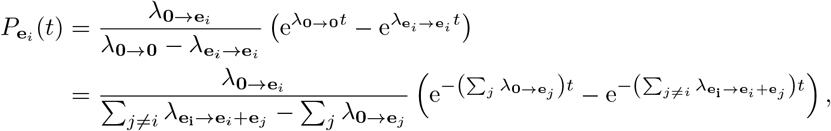

which agrees with Equation (5) for the case of *M* = 2.

## 3 Methods

### 3.1 Data sources

We obtained single-nucleotide variants (SNV) from 565 cases of lung adenocarcinoma from The Cancer Genome Atlas, and classified them into genes following human genome coordinates from the Ensembl Project [25]. All mutations were used to determine the baseline mutation rate. However, synonymous mutations were removed for the purpose of tallying prevalence of selected mutations because they are typically not selected. To estimate epistatic effects among four genes using the quadruplewise case *M* = 4, we restricted our analysis to the four genes most commonly fixed for mutations among the 565 samples.

### 3.2 Likelihood of observed frequencies of tumors

To quantify the flux associated with one mutation with no epistasis (Section 2.1), if we have a sample of *N* tumors within which *n* tumors exhibited the variant site, we can assume that the samples are taken at a similar time *T* from an initial state without the mutation assessed. The likelihood is binomial:

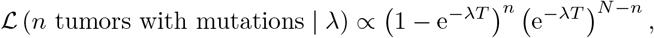

and can be maximized to obtain an estimate of *λ*.

With two mutations and accounting for epistatic effects (Section 2.2), the genotypes of *N* tumors can be subdivided into *n*_**0**_, *n*_*A*_, *n*_*B*_, and *n*_*AB*_, representing those tumors without any mutations, with only *A*, with only *B*, and with both *A* and *B*. To estimate all the fluxes, we set the time again at *t* = *T* and the likelihood is multinomial:

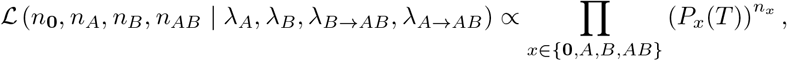

where *P*_*x*_(*T*) is given by Equations (2), (5), (6), and (7). The maximization of this likelihood will provide estimators of the fluxes.

For the general case with an arbitrary number of mutations (Section 2.3), a sample of *N* tumors that have been sequenced can be divided according to the somatic genotype in *S* attributed to each tumor. Let *n*_**x**_ be the total number of samples that have the somatic genotype **x** for each **x** ∈ *S*. We assume that the samples are taken at a similar time *T* (of potentially arbitrary unit) from an initial state without any of the *M* mutations assessed (i.e. the “normal” state **0**). Consequently, the probability *P*_**x**_(*T*) reflects the fraction of the cases observed with the somatic genotype **x**. The likelihood ℒ is then multinomial:

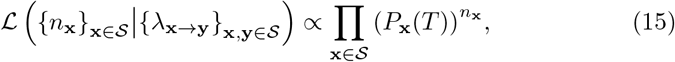

where 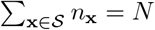, and *P*_**x**_(*T*) is dependent on the fluxes *λ*_**x**→**y**_ according to Equations (12) and (14).

We set *T* = 1 in Equation (15) to evaluate the likelihood, equating one unit to the average duration between the clonal origin of the tumor and tumor sampling; all fluxes derived with *T* = 1 are thus in units of 1 over this unit. To compute the integral on the right-hand side of the recursive formula in Equation (14), we used a trapezoidal rule with a resolution of 1,000 points between 0 and *T*. We tested higher resolutions, and our results were unchanged.

### 3.3 Estimation of parameters

We estimated the fluxes *λ*_**x**→**y**_ by maximizing the likelihood in Equation (15). We computed asymptotic confidence intervals for each of the flux estimates by computing the log-likelihood ratio and using Wilk’s theorem [26]. To validate model fit, we compared for each state **x** the probability *P*_**x**_(1) evaluated with the flux estimates Equation (14) to the observed fraction of the samples in each category. The values were equal for each somatic genotype.

To factor each flux into a mutation rate and a scaled selection coefficient, we assumed that the mutation rate per gene did not change with the acquisition of the somatic mutations of interest. To obtain gene-specific mutation rates for all possible variants of the gene, we used cancereffectsizeR [9]. cancereffectsizeR combines the gene-specific regression approach of dndsCV [27] with custom covariates to estimate gene-specific mutation rates. Underlying mutation rates for each gene were calculated based only on mutation rates of putative driver sites indicated by the presence of observed mutations in the gene in the cohort. Thus, the gene mutation rate *μ* was equal to the sum of the mutation rates for all observed variants of the gene, with the mutation rate for a variant *v* being equal to:

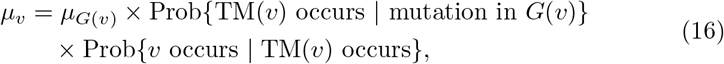

where *G*(*v*) is the gene of the variant *v, μ*_*G*(*v*)_ is the mutation rate for that gene for all possible variants (not only those observed) and TM(*v*) is the trinucleotide mutation of the variant *v*. To estimate the two probabilities in the right hand side of Equation (16) we used the proportion of the trinucleotide mutations across all data, and the number of times that the trinucleotide context appeared in the Genome Reference Consortium Human Build 38 [28], respectively.

## 4 Results

The genes that were most frequently mutated were tumor protein p53 (TP53, *n* = 243), low-density lipoprotein receptor-related protein 1B (LRP1B, *n* = 181), Kirsten rat sarcoma (KRAS, *n* = 150) and the tumor suppressor serine/threonine kinase 11 (STK11, *n* = 51). The numbers of patients with each somatic genotype informed the likelihood of our model, which provided estimates and confidence intervals for the flux and scaled selection coefficient of each mutation in the context of each somatic genotype for these four genes (Table 1).

**Table 1:** Fluxes, mutation rates, and scaled selection coefficients for a four-gene model of lung adenocarcinoma oncogenesis.

| Genotype | Mutation | Flux (CI)<sup>a</sup> | Mutation rate | SSC<sup>b</sup> (CI)<sup>a</sup> |
| --- | --- | --- | --- | --- |
| normal | *TP53* | 0.84 (0.73, 0.95) | 18.09e-8 | 4.62 (4.03, 5.27) |
| normal | *KRAS* | 0.31 (0.25, 0.38) | 1.99e-8 | 15.78 (12.78, 19.23) |
| normal | *STK11* | 0.08 (0.05, 0.11) | 2.54e-8 | 2.99 (1.93, 4.40) |
| normal | *LRP1B* | 0.19 (0.13, 0.26) | 15.60e-8 | 1.22 (0.86, 1.66) |
| *TP53* | *KRAS* | 0.29 (0.19, 0.41) | 1.99e-8 | 14.39 (9.50, 20.79) |
| *TP53* | *STK11* | 0.13 (0.07, 0.23) | 2.54e-8 | 5.21 (2.73, 8.86) |
| *TP53* | *LRP1B* | 0.84 (0.64, 1.07) | 15.60e-8 | 5.39 (4.13, 6.86) |
| *KRAS* | *TP53* | 0 (0, 0.30) | 18.09e-8 | 0 (0, 1.64) |
| *KRAS* | *STK11* | 0.82 (0.54, 1.19) | 2.54e-8 | 32.18 (21.43, 46.62) |
| *KRAS* | *LRP1B* | 0.56 (0.34, 0.86) | 15.60e-8 | 3.57 (2.18, 5.51) |
| *STK11* | *TP53* | 0 (0, 0.60) | 18.09e-8 | 0 (0, 3.34) |
| *STK11* | *KRAS* | 0 (0, 0.69) | 1.99e-8 | 0 (0, 34.62) |
| *STK11* | *LRP1B* | 0.41 (0.14, 0.91) | 15.60e-8 | 2.63 (0.92, 5.86) |
| *LRP1B* | *TP53* | 1.02 (0.51, 1.68) | 18.09e-8 | 5.65 (2.82, 9.28) |
| *LRP1B* | *KRAS* | 0 (0, 0.38) | 1.99e-8 | 0 (0, 18.91) |
| *LRP1B* | *STK11* | 0 (0, 0.23) | 2.54e-8 | 0 (0, 8.95) |
| *TP53+KRAS* | *STK11* | 0.52 (0.16, 1.21) | 2.54e-8 | 20.40 (6.48, 47.60) |
| *TP53+KRAS* | *LRP1B* | 0.21 (0, 1.17) | 15.60e-8 | 1.34 (0, 7.51) |
| *TP53+STK11* | *KRAS* | 0 (0, 1.21) | 1.99e-8 | 0 (0, 60.51) |
| *TP53+STK11* | *LRP1B* | 0.52 (0, 2.01) | 15.60e-8 | 3.34 (0, 12.88) |
| *KRAS+STK11* | *TP53* | 0 (0, 0.59) | 18.09e-8 | 0 (0, 3.26) |
| *KRAS+STK11* | *LRP1B* | 0.82 (0.35, 1.63) | 15.60e-8 | 5.28 (2.27, 10.47) |
| *TP53+LRP1B* | *KRAS* | 0.72 (0.45, 1.08) | 1.99e-8 | 36.03 (22.75, 53.96) |
| *TP53+LRP1B* | *STK11* | 0.21 (0.07, 0.42) | 2.54e-8 | 8.19 (2.75, 16.67) |
| *KRAS+LRP1B* | *TP53* | 0 (0, 1.00) | 18.09e-8 | 0 (0, 5.52) |
| *KRAS+LRP1B* | *STK11* | 0.08 (0, 0.85) | 2.54e-8 | 2.96 (0, 33.49) |
| *STK11+LRP1B* | *TP53* | 0 (0, 2.99) | 18.09e-8 | 0 (0, 16.55) |
| *STK11+LRP1B* | *KRAS* | 0 (0, 2.95) | 1.99e-8 | 0 (0, 148.07) |
| *TP53+KRAS+STK11* | *LRP1B* | 0.92 (0, 5.92) | 15.60e-8 | 5.90 (0, 37.97) |
| *TP53+KRAS+LRP1B* | *STK11* | 0.23 (0, 0.86) | 2.54e-8 | 9.06 (0, 33.98) |
| *TP53+STK11+LRP1B* | *KRAS* | 0 (0, 1.59) | 1.99e-8 | 0 (0, 80.04) |
| *KRAS+STK11+LRP1B* | *TP53* | 0 (0, 1.77) | 18.09e-8 | 0 (0, 9.76) |

<sup>a</sup> 95% confidence interval.
<sup>b</sup> Scaled selection coefficient (in millions).

Fixed-value estimates for the mutation rates for each gene over oncogenesis led to gene-specific linear relationships between the the scaled selection coefficient and the flux (Figure 1). A consequence of the multiplicative relationship between mutation rate and scaled selection coefficient is that mutation rate determined the slope of response of flux to scaled selection coefficient. Consequently, underlying mutation rate had a substantial effect on flux and prevalence of somatic genotypes, especially in the context of less-strongly selected mutations such as those in *LRP1B*. Mutations to *KRAS* were the least frequent of any gene (Table 1), and exhibited the lowest slope of flux in response to scaled selection coefficient.

**Figure 1:**
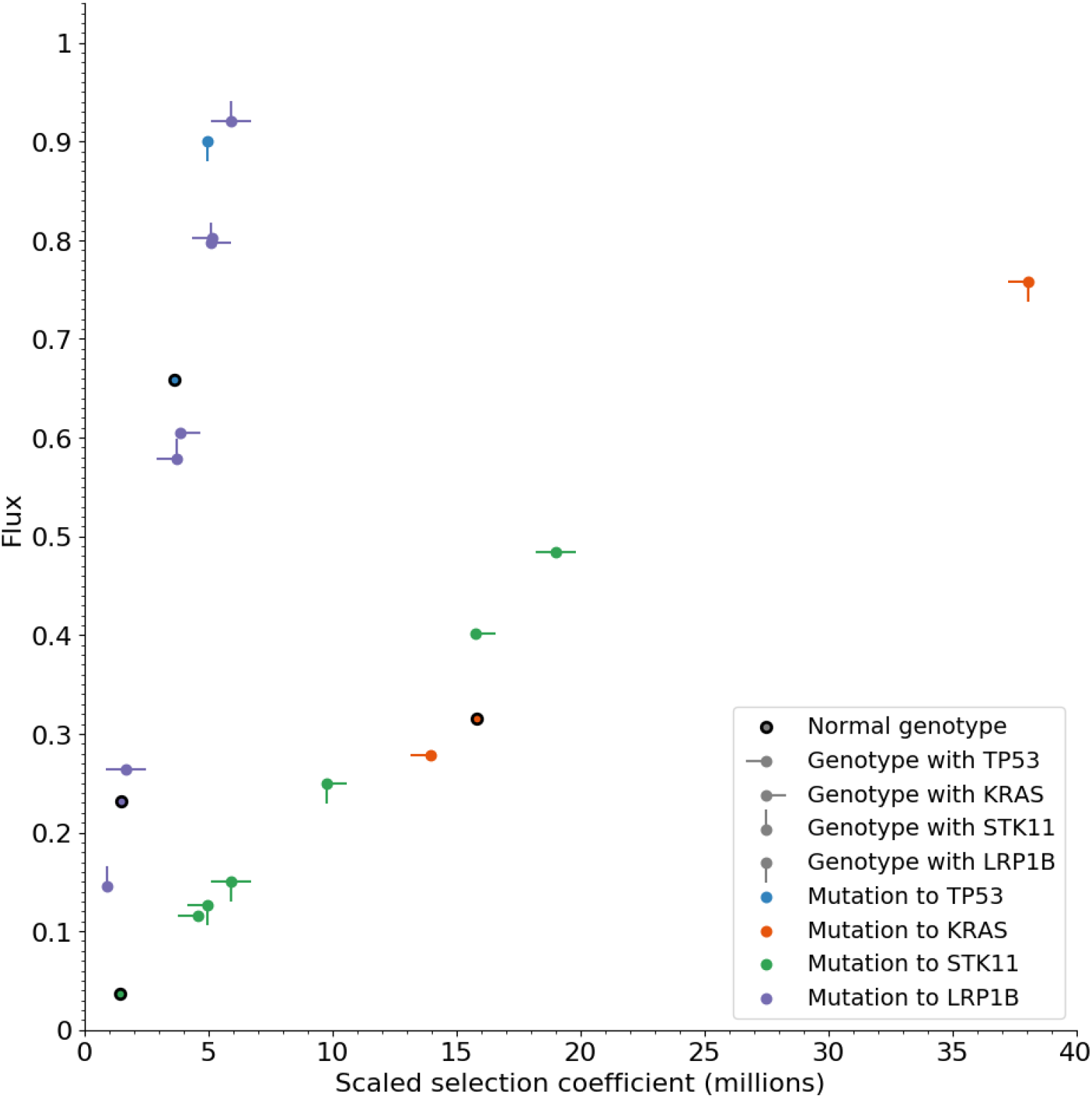
Estimates of positive scaled selection coefficients and positive genotypic fluxes for a four-gene model of lung adenocarcinoma oncogenesis. Each genotype (indicated by the directions of ticks superimposed on each point) has distinct effects on the flux that are mediated by epistasis affecting the scaled selection coefficients of new mutations (TP53, blue; KRAS, red; STK11, green; LRP1B, purple). No points are shown for any mutation whose estimated selection coefficient—and consequently the flux—was zero.

The slope of flux to scaled selection coefficient that is evident for somatic mutations of *KRAS* and *STK11* is lower than that for somatic mutations of *LRP1B* and *TP53*. However, the scaled selection coefficients for the positively selected mutations in *KRAS* and *STK11* are high for most tumor genotypes in lung adenocarcinoma (Figure 1). Scaled selection coefficients for *KRAS* and *TP53* mutations were especially high compared to *STK11* and LRPB1 at the initiation of oncogenesis. Consequently, the flux to *KRAS* from a normal somatic genotype was vastly higher than for any other mutation, with *TP53* a distant second, and *LRP1B* and *STK11* following. In later stages of oncogenesis, the flux to *LRP1B* can be relatively large. However, the larger magnitude of flux can clearly be attributed to its high mutation rate (Table 1, Figure 2), rather than to its relatively small scaled selection coefficient.

**Figure 2:**
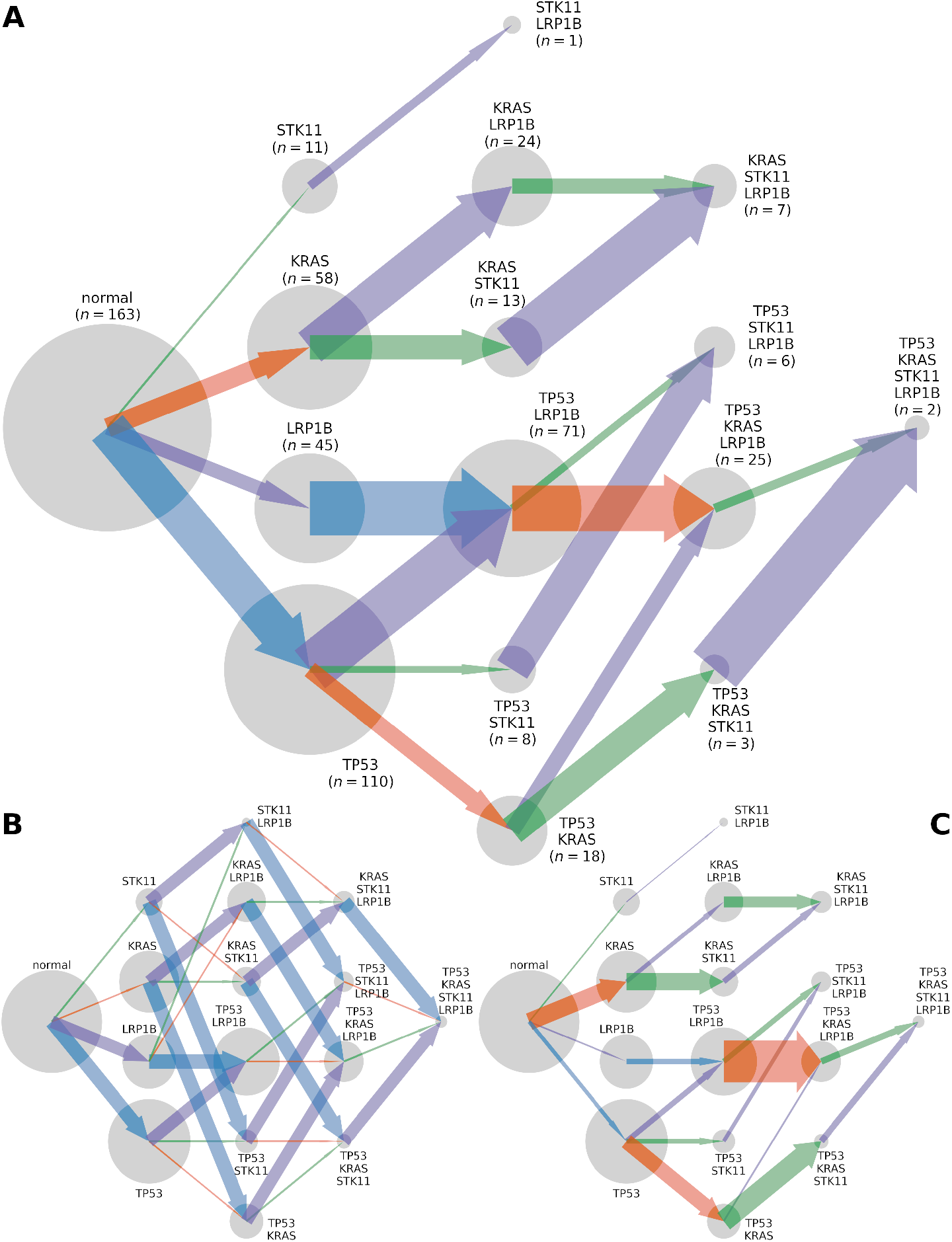
Trajectories of the somatic evolution by mutation of TP53, KRAS, LRP1B, and STK11, inferred from a total of 565 whole-exome sequenced lung adenocarcionma tumors. Genotypes (grey circles; areas are proportional to observed n for the genotype) evolve at (**A**) fluxes, (**B**) mutation rates, and (**C**) scaled selection coefficients that are proportional to the width of arrows pointing from one genotype to another, colored by the gene in which the mutation occurs (TP53, blue; KRAS, orange; STK11, green; LRP1B: purple).

Comparison of the scaled selection coefficient for fixation of the first mutations to the scaled selection coefficient for that mutation in non-normal genotypes frequently revealed an antagonistic epistatic effect of somatic mutations to oncogenic drivers (Table 1, Figure 2C). Antagonistic epistatic effects, in turn, likely explain the low number of patients that exhibit mutations in three or more of the four studied genes (Figure 2A). Antagonistic epistatic effects also partitioned the order of mutation fixation into three routes: (i) *STK11* then *LRP1B* (not very often); (ii) *KRAS*, then *STK11* or *LRP1B*, and then the remainder of *LRP1B* or (less often) of *STK11*; and (iii) more complex routes with a first fixation of either *LRP1B* or *TP53* (Figure 2A).

Some synergistic epistatic effects were also evident. For example, any genotype with a *KRAS* or *TP53* mutation substantially increased the scaled selection coefficient on *LRP1B* mutation compared to when neither gene was mutated (Table 1, Figure 2C). The largest significant synergistic epistatic effects arose in the presence of *TP53* and *LRP1B* when acquiring the *KRAS* mutation, in the presence of *KRAS* when acquiring *STK11*, and the presence of *TP53* and *KRAS* when acquiring the *STK11* mutation (Table 1, Figure 2C).

Examining the 24 possible paths of mutation fixation, it becomes clear that an initial *TP53* mutation promotes an increase in fitness of the second mutations and sometimes even the third mutation (Figure 3). A *KRAS* mutation from the normal genotype conveys substantial selective benefit. However, the selection coefficient decreases substantially for a second mutation in *TP53* or *LRP1B*, and remains about the same with a second mutation in *STK11* (Figure 3). Initial acquisition of *STK11* or *LRP1B* mutations rendered weak selection for additional mutations of the three other genes, with one notable exception: *TP53* following *LRP1B* (Figure 3).

**Figure 3:**
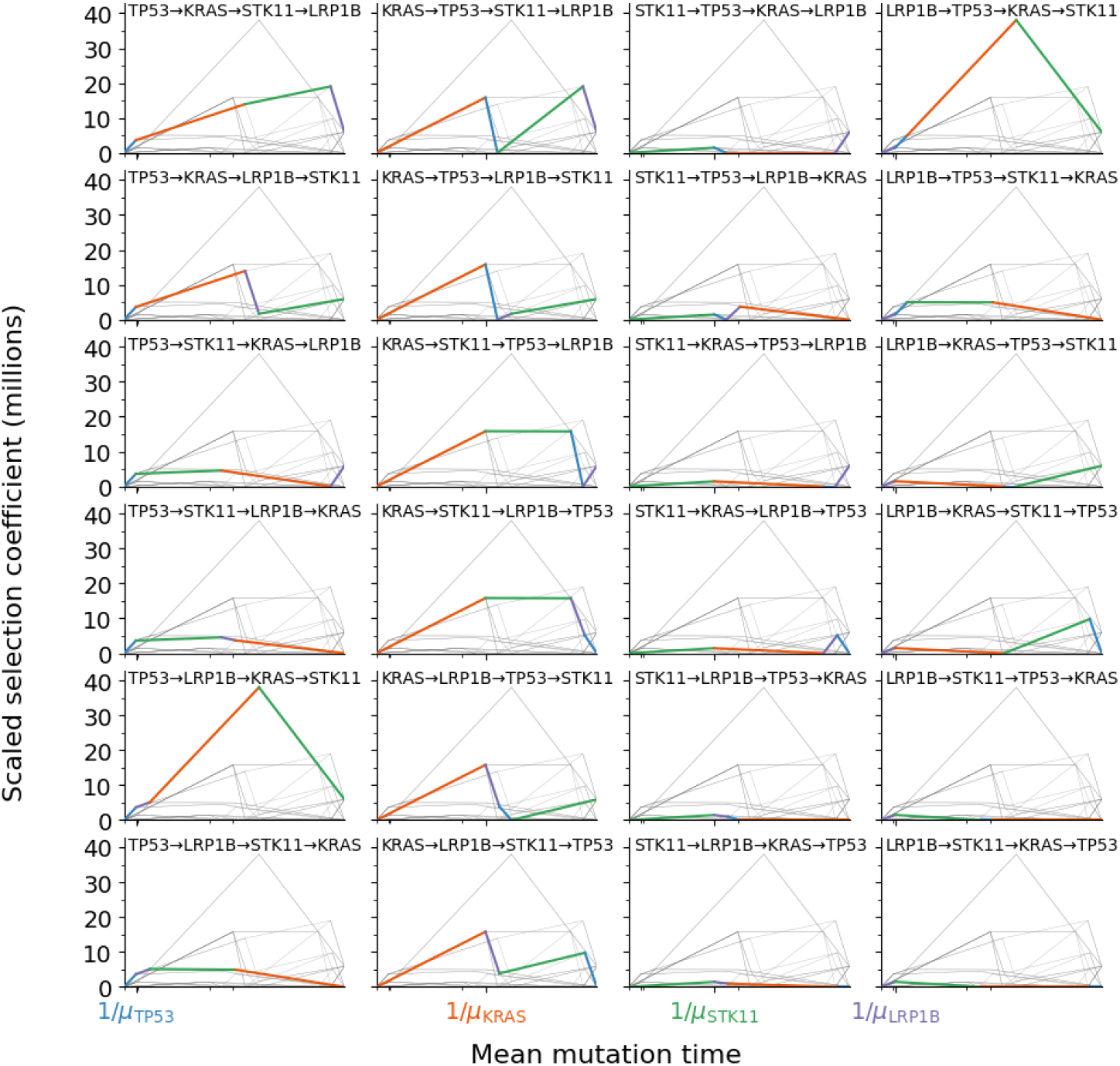
Mutation landscape depicting all paths of mutation acquisition in a fourgene model for lung adenocarcinoma. The four mutations incorporated are TP53 (blue, initial mutation in column one), KRAS (orange, initial mutation in column 2), STK11 (green, initial mutation in column three), and LRP1B (purple, initial mutation in column four). The mean time to acquire a mutation in a cancercompetent cell lineage (*x*-axis) quantifies how quickly each mutation will occur on a cellular level in the landscape of mutations (shorter is quicker), whereas the scaled selection coefficient (*y*-axis) is a measure of the benefit of the mutation to lineage proliferation and survival (the higher the selection coefficient, the more likely a mutation, once it occurs, will rapidly spread to high frequency in tumor tissue). In every subplot, one path is highlighted (colored curve, which corresponds to the order indicated by the gene names above the curve) and contrasted with all other possible paths (gray curves).

## 5 Discussion

Here we have shown how to estimate high-order epistatic effects on the somatic selection of cancer mutations. Our model enables computation of the flux in somatic genetic state of tissue from one genotype to another by single mutations. Application of our model to 565 lung adenocarcinoma samples pro-vided estimates of 32 fluxes to the 4 most commonly mutated genes (*TP53, LRP1B, KRAS* and *STK11*), and showed that the flux to each of those four genes depended on the current somatic genotype. Many genotypes exhibited an antagonistic epistatic effect that resulted in zero or near-zero fluxes out of that genotype. Antagonistic epistatic effects likely partly explain the relatively low number of tumor samples that contain mutations in three or more of these commonly-mutated genes.

By estimating the neutral mutation rates of each gene, we enabled computation of the corresponding scaled selection coefficients that quantify the degree to which the mutations increased survival and proliferation. We found that *KRAS* mutations exhibited scaled selection coefficients that were especially large and were a major reason for their high fluxes, especially from the *TP53* +*LRP1B* genotype. In contrast, most of the flux to *LRP1B* mutations—and thus the large number of samples with *LRP1B* mutations—can be explained by *LRP1B* ‘s large mutation rate, not by a large cancer effect. Low effect size indicates that abrogating the effect of these *LRP1B* mutations will have modest impact on tumor cell growth and survival. However, *LRP1B* mutations could be highly relevant to prognosis and therapy in other ways. For instance, lung adenocarcinoma tumor mutations in *LRP1B* have been associated with chronic obstruction pulmonary disease in patients [29], and have also been suggested to be useful predictors of response to immune checkpoint inhibitors [30]. Moreover, our analyses of epistatic interactions corroborates previous studies that suggest that *LRP1B* cooperates with *TP53* to induce strong selection for high-effect driver mutations of *KRAS* [5]. An intermediate mutation of *LRP1B* might compose a valley in the cancer fitness landscape that needs to be crossed to obtain a substantial advantage in selection with a final mutation in *KRAS* [31].

*KRAS* and *TP53* have been suggested to play a key role in the initial stages of the somatic evolution of many lung adenocarcinomas [32–34]. Our analysis provided a result consistent with these findings: *KRAS* is subject to a synergistic epistatic effect on its selection in the context of *TP53* mutations. Interestingly, our quantification of these epistatic effects argues that the order in which *KRAS* and *TP53* /*STK11* mutations are acquired is relevant to their selective effect. Acquisition of a *TP53* mutation before a *KRAS* mutation does little to change selection on a new *KRAS* mutation. However, if a *KRAS* mutation is acquired first, selection on *TP53* mutation almost disappears. Conversely, acquisition of a *STK11* mutation from a normal state eliminates strong selection on *KRAS* —but if *KRAS* is acquired first, there is strong selection for *STK11. TP53* and *STK11* have been identified as determinants of distinct subsets of lung adenocarcinomas dominated by *KRAS* mutants [35]. Importantly, patients whose tumors have *KRAS* driver mutations but no *TP53* driver mutations have better overall survival than those with both, especially in the absence of *STK11* mutations [36, 37]. Indeed—according to our results— the better prognosis of *KRAS* without *TP53* or *STK11* arises more often when a *LRP1B* mutation occurs immediately after a *KRAS* mutation initiates driver-based divergence from the normal genotype.

*STK11* has previously been identified as a mutation that occurs relatively early in the oncogenesis of lung adenocarcinoma [34]. Our results indicate that an early, relatively simple route of mutations starts with a *STK11* mutation and continues with an *LRP1B* mutation, without further mutation of *KRAS* or *TP53* prior to tumor resection, but it does not occur very often. We have estimated strong selection for *STK11* mutations after *KRAS* mutation, indicating that if *KRAS* mutations occur early, then *STK11* mutations would be strongly selected, and once mutated, would likely reach high tumor frequencies not long after. However, we have also found a relatively constant selection for *STK11* from other genotypes, that—despite being lower than when *KRAS* mutates first—suggests that *STK11* does not arise exclusively early during tumor initiation.

We focused here on single-nucleotide variants that affect the genes studied. However, there are other somatic factors that could have epistatic effects, such as copy-number alterations and structural variants. Our model could incorporate those factors and compute the fluxes associated with their fixation. However, the estimation of the underlying mutation rate for copy number alterations and structural variants remains a challenge. Consequently, quantifying the strength of selection operating on them is not presently feasible. Thus far, research has indicated that copy-number changes and structural variants do not substantially influence selection on simple nucleotide variations, and have instead orthogonal effects in cancers [38].

Confidence intervals for our estimates of epistatic selection became wider as the genotypes included mutations in more genes, because the number of tumors with mutations in three or four of these commonly-mutated genes tended to be very low. To reduce the parameter uncertainty, future research should explore larger data sets by aggregating whole-exome sequences and panel data from multiple sources. Alternatively, data that includes tumor samples taken at multiple time points during tumor evolution could be incorporated to directly inform the order at which mutations occur, reducing uncertainty. When considering therapy, incorporation of multiple samples from the same tumor can play an important role, because genetically heterogeneous tumors can exert proclivity toward very different evolutionary outcomes [39]. Very soon after therapy, regions with clones resistant to therapy can become widespread affecting the prognosis for the patient [15, 40]. The equations we have derived enable estimation of the time-dependent probability of each somatic genotype given the flux values, and could be modified to incorporate changes in mutation and selection after therapy. Thus, our theory could be extended to applications quantifying cancer effects on tumor samples at metachronous time points.

Using an established approach [9], we were able to estimate the mutation rate in each gene. However, this approach does not evaluate whether somatic genotypes vary in gene mutation rate. Mutation rates that depend on somatic genotype could affect the co-occurence or mutual exclusivity of future mutations [40–43]. For example, it has long been suggested that mutations in *TP53* impair response to DNA damage, which would typically lead to higher gene-mutation rates thenceforward and could even increase mutation rates in specific genomic hotspots [40, 44].

Environmental exposures including carcinogens and chemotherapy also have the potential to change during the course of tumor evolution and affect mutation rates [40, 45, 46]. For instance, tobacco smoking is a major factor that increases the mutation rate of certain oncogenic sites in lung cancer, including *KRAS* [10, 47]. Incorporation of differences in mutation rates depending on endogenous and exogenous mutational processes associated with somatic genotype and environmental exposures such as smoking, could enable increasingly precise estimation of selection coefficients attributable to each mutation.

In summary, we have derived methods enabling estimation of the fluxes and selection coefficients dependent on genotypes among somatic cancer mutations, and have employed them to obtain epistatic effects among the four most commonly mutated genes in lung adenocarcinoma. We revealed several strong antagonistic and synergistic epistatic effects that reduce the space of possible oncogenic trajectories, and quantified the likelihoods of tumors evolving along each path. Determining these most likely trajectories can help in future applications to determine optimal personalized therapies that account for the current somatic genotype of tumor tissue from a patient, and that are designed to forestall specific somatic genotypic trajectories that are most likely to be forthcoming.

## 6 Acknowledgements

Data used for this research were generated by the TCGA Research Network: https://www.cancer.gov/tcga. We would like to thank Katherine Brumberg and Vincent Cannataro for early discussions of ideas regarding this research, and Krishna Dasari for discussions in the later stages of the manuscript and for contributing code to obtain the mutation rate per variant. This research was supported by NSF IOS 1934848, NIH 1R01LM013385, and NIH NIDCR 1P50DE030707.

## References

[1] Xiaoyue Wang, Audrey Q Fu, Megan E McNerney, and Kevin P White. Widespread genetic epistasis among cancer genes. Nature communications, 5(1):1–10, 2014.

[2] Trachette Jackson, Natalia Komarova, and Kristin Swanson. Mathematical oncology: using mathematics to enable cancer discoveries. The American Mathematical Monthly, 121(9):840–856, 2014.

[3] R. Manavalan and S. Priya. Genetic interactions effects for cancer disease identification using computational models: a review. Medical & Biological Engineering & Computing, 59(4):733–758, 2021.

[4] Kieran Elmes, Fabian Schmich, Ewa Szczurek, Jeremy Jenkins, Niko Beerenwinkel, and Alex Gavryushkin. Learning epistatic gene interactions from perturbation screens. Plos one, 16(7):e0254491, 2021.

[5] Michael I Klein, Vincent L Cannataro, Jeffrey P Townsend, Scott Newman, David F Stern, and Hongyu Zhao. Identifying modules of cooperating cancer drivers. Molecular systems biology, 17(3):e9810, 2021.

[6] Daniel M Weinreich, Yinghong Lan, Jacob Jaffe, and Robert B Heckendorn. The influence of higher-order epistasis on biological fitness landscape topography. Journal of statistical physics, 172(1):208–225, 2018.

[7] Daniel M Weinreich, Yinghong Lan, C Scott Wylie, and Robert B Heckendorn. Should evolutionary geneticists worry about higher-order epistasis? Current opinion in genetics & development, 23(6):700–707, 2013.

[8] Anastasia Baryshnikova, Michael Costanzo, Chad L Myers, Brenda Andrews, and Charles Boone. Genetic interaction networks: toward an understanding of heritability. Annual review of genomics and human genetics, 14:111–133, 2013.

[9] Vincent L Cannataro, Stephen G Gaffney, and Jeffrey P Townsend. Effect sizes of somatic mutations in cancer. JNCI: Journal of the National Cancer Institute, 110(11):1171–1177, 2018.

[10] Chichun Tan, Jeffrey D Mandell, Krishna Dasari, Vincent L Cannataro, Jorge A Alfaro-Murillo, and Jeffrey P Townsend. Heavy mutagenesis by tobacco leads to lung adenocarcinoma tumors with kras g12 mutations other than g12d, leading kras g12d tumors-on average-to exhibit a lower mutation burden. Lung Cancer, 166:265–269, 2021.

[11] Navodit Misra, Ewa Szczurek, and Martin Vingron. Inferring the paths of somatic evolution in cancer. Bioinformatics, 30(17):2456–2463, 2014.

[12] James RM Black and Nicholas McGranahan. Genetic and non-genetic clonal diversity in cancer evolution. Nature Reviews Cancer, 21(6):379–392, 2021.

[13] Jon F Wilkins, Vincent L Cannataro, Brian Shuch, and Jeffrey P Townsend. Analysis of mutation, selection, and epistasis: an informed approach to cancer clinical trials. Oncotarget, 9(32):22243, 2018.

[14] Krishna Dasari, Jason A Somarelli, Sudhir Kumar, and Jeffrey P Townsend. The somatic molecular evolution of cancer: Mutation, selection, and epistasis. Progress in biophysics and molecular biology, 165:56–65, 2021.

[15] Russell C Rockne, Andrea Hawkins-Daarud, Kristin R Swanson, James P Sluka, James A Glazier, Paul Macklin, David A Hormuth, Angela M Jarrett, Ernesto A B F Lima, J Tinsley Oden, George Biros, Thomas E Yankeelov, Kit Curtius, Ibrahim Al Bakir, Dominik Wodarz, Natalia Komarova, Luis Aparicio, Mykola Bordyuh, Raul Rabadan, Stacey D Finley, Heiko Enderling, Jimmy Caudell, Eduardo G Moros, Alexander R A Anderson, Robert A Gatenby, Artem Kaznatcheev, Peter Jeavons, Nikhil Krishnan, Julia Pelesko, Raoul R Wadhwa, Nara Yoon, Daniel Nichol, Andriy Marusyk, Michael Hinczewski, and Jacob G Scott. The 2019 mathematical oncology roadmap. Physical Biology, 16(4):041005, 2019.

[16] Brett C Bade and Charles S Dela Cruz. Lung cancer 2020: epidemiology, etiology, and prevention. Clinics in chest medicine, 41(1):1–24, 2020.

[17] U.S. Cancer Statistics Working Group. U.S. Cancer Statistics Data Visualizations Tool, based on 2020 submission data (1999-2018). U.S. Department of Health and Human Services, Centers for Disease Control and Prevention and National Cancer Institute; http://www.cdc.gov/cancer/dataviz, released in June 2021, 2021.

[18] David J Myers. Cancer, lung adenocarcinoma. StatPearls Publishing; https://www.statpearls.com/articlelibrary/viewarticle/24486/, last updated when retrieved, September 2021, 2021.

[19] Cancer Genome Atlas Research Network. Comprehensive molecular profiling of lung adenocarcinoma: The Cancer Genome Atlas research network. Nature, 511(7511):543–550, 2014.

[20] John H Gillespie. Some properties of finite populations experiencing strong selection and weak mutation. The American Naturalist, 121(5):691–708, 1983.

[21] David M Roy, Logan A Walsh, and Timothy A Chan. Driver mutations of cancer epigenomes. Protein & cell, 5(4):265–296, 2014.

[22] Joris van de Haar, Sander Canisius, K Yu Michael, Emile E Voest, Lodewyk FA Wessels, and Trey Ideker. Identifying epistasis in cancer genomes: a delicate affair. Cell, 177(6):1375–1383, 2019.

[23] Andrei Kolmogoroff. Über die analytischen Methoden in der Wahrschein-lichkeitsrechnung. Mathematische Annalen, 104(1):415–458, 1931.

[24] L. Perko. Differential Equations and Dynamical Systems. Springer, third edition, 2001.

[25] Kevin L Howe, Premanand Achuthan, James Allen, Jamie Allen, Jorge Alvarez-Jarreta, M Ridwan Amode, Irina M Armean, Andrey G Azov, Ruth Bennett, Jyothish Bhai, Konstantinos Billis, Sanjay Boddu, Mehrnaz Charkhchi, Carla Cummins, Luca Da Rin Fioretto, Claire Davidson, Kamalkumar Dodiya, Bilal El Houdaigui, Reham Fatima, Astrid Gall, Carlos Garcia Giron, Tiago Grego, Cristina Guijarro-Clarke, Leanne Haggerty, Anmol Hemrom, Thibaut Hourlier, Osagie G Izuogu, Thomas Juettemann, Vinay Kaikala, Mike Kay, Ilias Lavidas, Tuan Le, Diana Lemos, Jose Gonzalez Martinez, José Carlos Marugán, Thomas Maurel, Aoife C McMahon, Shamika Mohanan, Benjamin Moore, Matthieu Muffato, Denye N Oheh, Dimitrios Paraschas, Anne Parker, Andrew Parton, Irina Prosovetskaia, Manoj P Sakthivel, Ahamed I Abdul Salam, Bianca M Schmitt, Helen Schuilenburg, Dan Sheppard, Emily Steed, Michal Szpak, Marek Szuba, Kieron Taylor, Anja Thormann, Glen Threadgold, Brandon Walts, Andrea Winterbottom, Marc Chakiachvili, Ameya Chaubal, Nishadi De Silva, Bethany Flint, Adam Frankish, Sarah E Hunt, Garth R IIsley, Nick Langridge, Jane E Loveland, Fergal J Martin, Jonathan M Mudge, Joanella Morales, Emily Perry, Magali Ruffier, John Tate, David Thybert, Stephen J Trevanion, Fiona Cunningham, Andrew D Yates, Daniel R Zerbino, and Paul Flicek. Ensembl 2021. Nucleic Acids Research, 49(D1):D884–D891, 11 2020.

[26] Samuel S Wilks. The large-sample distribution of the likelihood ratio for testing composite hypotheses. The annals of mathematical statistics, 9(1):60–62, 1938.

[27] Iñigo Martincorena, Keiran M Raine, Moritz Gerstung, Kevin J Dawson, Kerstin Haase, Peter Van Loo, Helen Davies, Michael R Stratton, and Peter J Campbell. Universal patterns of selection in cancer and somatic tissues. Cell, 171(5):1029–1041, 2017.

[28] The Genome Reference Consortium. Genome Reference Consortium Human Build 38. https://www.ncbi.nlm.nih.gov/grc, 2021.

[29] Dakai Xiao, Fuqiang Li, Hui Pan, Han Liang, Kui Wu, and Jianxing He. Integrative analysis of genomic sequencing data reveals higher prevalence of LRP1B mutations in lung adenocarcinoma patients with COPD. Scientific reports, 7(1):1–8, 2017.

[30] Shaowei Lan, Hui Li, Ying Liu, Lixia Ma, Xianhong Liu, Yan Liu, Shi Yan, and Ying Cheng. Somatic mutation of LRP1B is associated with tumor mutational burden in patients with lung cancer. Lung cancer, 132:154–156, 2019.

[31] Natalia L Komarova, Leili Shahriyari, and Dominik Wodarz. Complex role of space in the crossing of fitness valleys by asexual populations. Journal of The Royal Society Interface, 11(95):20140014, 2014.

[32] Maria Grzes, Magdalena Oron, Zuzanna Staszczak, Akanksha Jaiswar, Magdalena Nowak-Niezgoda, and Dawid Walerych. A driver never works alone—interplay networks of mutant p53, MYC, RAS, and other universal oncogenic drivers in human cancer. Cancers, 12(6):1532, 2020.

[33] Roy S Herbst, Daniel Morgensztern, and Chris Boshoff. The biology and management of non-small cell lung cancer. Nature, 553(7689):446–454, 2018.

[34] Moritz Gerstung, Clemency Jolly, Ignaty Leshchiner, Stefan C Dentro, Santiago Gonzalez, Daniel Rosebrock, Thomas J Mitchell, Yulia Rubanova, Pavana Anur, Kaixian Yu, Maxime Tarabichi, Amit Deshwar, Jeff Wintersinger, Kortine Kleinheinz and Ignacio Vázquez-García, Kerstin Haase, Lara Jerman, Subhajit Sengupta, Geoff Macintyre, Salem Malikic, Nilgun Donmez, Dimitri G Livitz, Marek Cmero, Jonas Demeulemeester, Steven Schumacher, Yu Fan, Xiaotong Yao, Juhee Lee, Matthias Schlesner, Paul C Boutros, David D Bowtell, Hongtu Zhu, Gad Getz, Marcin Imielinski, Rameen Beroukhim, S Cenk Sahinalp, Yuan Ji, Martin Peifer, Florian Markowetz, Ville Mustonen, Ke Yuan, Wenyi Wang, Quaid D Morris, PCAWG Evolution & Heterogeneity Working Group, Paul T Spellman, David C Wedge, Peter Van Looand, and PCAWG Consortium. The evolutionary history of 2,658 cancers. Nature, 578(7793):122–128, 2020.

[35] Ferdinandos Skoulidis, Lauren A Byers, Lixia Diao, Vassiliki A Papadimitrakopoulou, Pan Tong, Julie Izzo, Carmen Behrens, Humam Kadara, Edwin R Parra, Jaime Rodriguez Canales, Jianjun Zhang, Uma Giri, Jayanthi Gudikote, Maria A. Cortez, Chao Yang, Youhong Fan, Michael Peyton, Luc Girard, Kevin R. Coombes, Carlo Toniatti, Timothy P. Heffernan, Murim Choi, Garrett M. Frampton, Vincent Miller, John N. Weinstein, Roy S. Herbst, Kwok-Kin Wong, Jianhua Zhang, Padmanee Sharma, Gordon B. Mills, Waun K. Hong, John D. Minna, James P. Allison, Andrew Futreal, Jing Wang, Ignacio I. Wistuba, and John V. Heymach. Co-occurring genomic alterations define major subsets of KRAS-mutant lung adenocarcinoma with distinct biology, immune profiles, and therapeutic vulnerabilities. Cancer discovery, 5(8):860–877, 2015.

[36] Linnéa La Fleur, Elin Falk-Sörqvist, Patrik Smeds, Anders Berglund, Magnus Sundström, Johanna SM Mattsson, Eva Brandén, Hirsh Koyi, Johan Isaksson, Hans Brunnström, Mats Nilsson, Patrick Micke, Lotte Moens, and Johan Botling. Mutation patterns in a population-based non-small cell lung cancer cohort and prognostic impact of concomitant mutations in KRAS and TP53 or STK11. Lung Cancer, 130:50–58, 2019.

[37] Nicolas Pécuchet, Pierre Laurent-Puig, Audrey Mansuet-Lupo, Antoine Legras, Marco Alifano, Karine Pallier, Audrey Didelot, Laure Gibault, Claire Danel, Pierre-Alexandre Just, Marc Riquet, Francoise Le Pimpec-Barthes, Diane Damotte, Elisabeth Fabre, and Hélène Blons. Different prognostic impact of STK11 mutations in non-squamous non-small-cell lung cancer. Oncotarget, 8(14):23831, 2017.

[38] Yifeng Tao, Ashok Rajaraman, Xiaoyue Cui, Ziyi Cui, Haoran Chen, Yuanqi Zhao, Jesse Eaton, Hannah Kim, Jian Ma, and Russell Schwartz. Assessing the contribution of tumor mutational phenotypes to cancer progression risk. PLoS computational biology, 17(3):e1008777, 2021.

[39] Suzan Farhang-Sardroodi, Amir H Darooneh, Mohammad Kohandel, and Natalia L Komarova. Environmental spatial and temporal variability and its role in non-favoured mutant dynamics. Journal of The Royal Society Interface, 16(157):20180781, 2019.

[40] J Nicholas Fisk, Amandeep R Mahal, Alex Dornburg, Stephen G Gaffney, Sanjay Aneja, Joseph N Contessa, David Rimm, B Yu James, and Jeffrey P Townsend. Premetastatic shifts of endogenous and exogenous mutational processes support consolidative therapy in EGFR-driven lung adenocarcinoma. Cancer letters, 526(1):346–351, 2022.

[41] Ahrim Youn and Richard Simon. Using passenger mutations to estimate the timing of driver mutations and identify mutator alterations. BMC Bioinformatics, 14(1):1–11, 2013.

[42] Edward J Fox, Marc J Prindle, and Lawrence A Loeb. Do mutator mutations fuel tumorigenesis? Cancer and Metastasis Reviews, 32(3):353–361, 2013.

[43] Keiichi Hatakeyama, Keiichi Ohshima, Takeshi Nagashima, Shumpei Ohnami, Sumiko Ohnami, Masakuni Serizawa, Yuji Shimoda, Koji Maruyama, Yasuto Akiyama, Kenichi Urakami, Masatoshi Kusuhara, Tohru Mochizuki, and Ken Yamaguchi. Molecular profiling and sequential somatic mutation shift in hypermutator tumours harbouring POLE mutations. Scientific reports, 8(1):1–12, 2018.

[44] Ian PM Tomlinson, MR Novelli, and WF Bodmer. The mutation rate and cancer. Proceedings of the National Academy of Sciences, 93(25):14800–14803, 1996.

[45] Vincent L. Cannataro, Jeffrey D. Mandell, and Jeffrey P. Townsend. Attribution of cancer origins to endogenous, exogenous, and preventable mutational processes. Molecular Biology and Evolution, page msac084, 2022.

[46] Natalia L Komarova and Dominik Wodarz. Evolutionary dynamics of mutator phenotypes in cancer: implications for chemotherapy. Cancer Research, 63(20):6635–6642, 2003.

[47] H Kadara, M Choi, J Zhang, ER Parra, J Rodriguez-Canales, SG Gaffney, Z Zhao, C Behrens, J Fujimoto, C Chow, Y Yoo, N Kalhor, C Moran, D Rimm, S Swisher, DL Gibbons, J Heymach, E Kaftan, JP Townsend, TJ Lynch, J Schlessinger,, J Lee, RP Lifton, II Wistuba, and RS Herbst. Whole-exome sequencing and immune profiling of early-stage lung adenocarcinoma with fully annotated clinical follow-up. Annals of Oncology, 28(1):75–82, 2017.

